# Transcriptomic and metabolomic responses to carbon and nitrogen sources in *Methylomicrobium album BG8*

**DOI:** 10.1101/2021.02.18.431927

**Authors:** Scott Sugden, Marina Lazic, Dominic Sauvageau, Lisa Y. Stein

## Abstract

Methanotrophs use methane as their sole carbon and energy source and represent an attractive platform for converting single-carbon feedstocks into value-added compounds. Optimizing these species for biotechnological applications involves choosing an optimal growth substrate based on an understanding of cellular responses to different nutrients. Although many studies of methanotrophs have examined growth rate, yield, and central carbon flux in cultures grown with different carbon and nitrogen sources, few studies have examined more global cellular responses to different media. Here, we evaluated global transcriptomic and metabolomic profiles of *Methylomicrobium album* BG8 when grown with methane or methanol as the carbon source and nitrate or ammonium as the nitrogen source. We identified five key physiological changes during growth on methanol: *M. album* BG8 cultures upregulated transcripts for the Entner-Doudoroff and pentose phosphate pathways for sugar catabolism, produced more ribosomes, remodeled its phospholipid membrane, activated various stress response systems, and upregulated glutathione-dependent formaldehyde detoxification. When using ammonium, *M. album* BG8 upregulated *haoAB* hydroxylamine dehydrogenase and the overall central metabolic activity; whereas when using nitrate, cultures upregulated genes for nitrate assimilation and conversion. Overall, we identified several nutrient source-specific responses that could provide a valuable basis for future research on the biotechnological optimization of these species.

**IMPORTANCE:** Methanotrophs are gaining increasing interest for their biotechnological potential to convert single-carbon compounds into value-added products such as industrial chemicals, fuels, and bioplastics. Optimizing these species for biotechnological applications requires a detailed understanding of how cellular activity and metabolism varies across different growth substrates. Although each of the two most commonly used carbon sources (methane or methanol) and nitrogen sources (ammonium or nitrate) in methanotroph growth media have well-described advantages and disadvantages in an industrial context, their effects on global cellular activity remain poorly characterized. Here, we comprehensively describe the transcriptomic and metabolomic changes that characterize the growth of an industrially promising methanotroph strain on multiple combinations of carbon and nitrogen sources. Our results represent a more holistic evaluation of cellular activity than previous studies of core metabolic pathways and provide a valuable basis for the future biotechnological optimization of these species.

## INTRODUCTION

Methanotrophs are a taxonomically diverse group of bacteria capable of using methane (CH_4_) and other one-carbon molecules as their sole source of energy and carbon (1). All aerobic methanotrophs share the same initial steps of their methane utilization pathway: methane is oxidized to methanol by methane monooxygenase (MMO) and methanol is then converted to formaldehyde by methanol dehydrogenase (MDH) (1). The metabolic pathways downstream from formaldehyde vary among species and growth conditions; formaldehyde can either be incorporated into cell biomass via the serine cycle in alphaproteobacterial methanotrophs or the ribulose monophosphate (RuMP) cycle in gammaproteobacterial methanotrophs, or it is further oxidized to formate using either the tetrahydromethanopterin pathway or formaldehyde dehydrogenase (2). Formate is then oxidized to CO_2_ by formate dehydrogenase. In gammaproteobacterial methanotrophs, the RuMP pathway feeds fructose-6-phosphate into the Embden-Meyerhof-Parnas (EMP), Entner-Doudoroff (ED), or pentose phosphate (PP) pathways (2).

Although methane is considered the primary carbon substrate for methanotrophs, many culture-based studies addressing the biotechnological applications of these species have explored the use of alternative growth substrates such as methanol (3–6). In some cases, alternative carbon sources may be desirable because they are cost-efficient, easier to upscale, or are the target of bioremediation programs (7). Alternative growth sources may also affect growth rates or increase the yield of value-added compounds, which are the target of ongoing bioindustrial research (4, 8). Methanol has been one of the most widely studied carbon sources aside from methane because it is easier to use in continuous culture systems (9), can be inexpensively synthesized from CO_2_ or CH_4_ (10), and avoids gas-liquid mass transfer issues associated with insoluble gases such as methane (11).

Many alphaproteobacterial methanotrophs exhibit a longer lag phase, slower growth rate, and lower final yield when grown on methanol, but several gammaproteobacterial methanotrophs exhibit robust growth on this substrate (3, 12, 13). Previous studies of gammaproteobacterial growth in methanol have largely focused on growth rate, yield, or flux through central carbon pathways (3, 5, 6, 14–16). For example, both *Methylotuvimicrobium alcaliphilum 20Z* and *Methylomonas* sp. DH-1 were shown to upregulate the EMP pathway during growth on methanol (5, 17), whereas *Methylotuvimicrobium buryantese* 5GB1 favored the ED pathway, which appears to be essential to this species (18), along with an incomplete TCA cycle (6). Because most of the electrons required by MMO are provided by the MDH co-factor pyrroloquinoline-quinone (PQQ) (11), growth on methanol has been proposed to free MDH-derived electrons for use in ATP production, though this hypothesis has not been widely tested outside of *M. buryatense* 5GB1 (6). Despite the value of these targeted carbon flux studies, there is limited information on global cellular responses and adaptations of methanotrophs to growth on methanol outside of changes in their central carbon pathways.

Nitrogen source is another important consideration for the growth of methanotrophs as it also has the potential to influence growth rate and other aspects of cellular physiology. Methanotrophs possess a diverse suite of genes for importing and metabolizing nitrogen, and some are capable of nitrification or denitrification activities (19–21). Ammonium may be a more bioenergetically favorable nitrogen source because it can be directly incorporated into cell biomass, but it can also be oxidized by MMO, competitively inhibiting methane oxidation and causing the formation of toxic intermediates like hydroxylamine (22). Because of competitive inhibition by ammonium, nitrate has been used in the typical growth medium for culture-based studies of methanotrophs (3), even though some species grow better on ammonium and are able to detoxify hydroxylamine (3, 22, 23).

Recent studies of methanotrophic growth have begun to identify strain-specific preferences based on different combinations of carbon-nitrogen sources that also affect metabolite pools (3–5, 13, 23). These studies have also shown that the effects of methanol toxicity exceed the effects of nitrogen source for cultures grown in methane. In this study, we used global transcriptomic and metabolomic analyses to holistically evaluate the metabolism and physiology of *M. album* BG8 grown with different combinations of carbon-nitrogen sources. *M. album* BG8 was recently classified into a separate genus from *M. buryatense* and *M. alcaliphilum* (24), grows to its highest optical density on methanol rather than methane (3), and prefers growth on nitrate rather than ammonium (3, 23). We first evaluated the strategies adopted by *M. album* BG8 to maintain equal growth rates in either methane or methanol, and assessed whether these strategies differ depending on whether the culture is provided with nitrate or ammonium. To determine if methanol alters nitrogen source preference, we additionally compared the nitrogen source response in methanol to the nitrogen source response to methane. Our results provide information on how *M. album* BG8 responds to different nutrient environments and serve as a valuable context for optimizing its growth and facilitating its industrialization towards the production of value-added products using single carbon compounds as feedstocks.

## RESULTS

We harvested *M. album* BG8 cultures grown in four different growth conditions derived from the combinations of two carbon sources (methane or methanol) and two nitrogen sources (ammonium or nitrate). All cultures exhibited similar growth rates and yields, regardless of the carbon-nitrogen source combination (**Fig. 1**). RNA-seq analysis from the cultures yielded 176,337,841 reads across 12 samples (triplicate of each carbon-nitrogen combination), representing 3,772 of the 3,794 annotated sequences in the published *M. album* BG8 genome (**Table S1**). Of these genes, 567 transcripts comprising 7.81% of all reads were not annotated in the Clusters of Orthologous Groups (COG) database, leaving 3,205 meaningfully classified genes that were used for downstream over-representation analyses. Genes for particulate methane monooxygenase (*pmoCAB*) were among the most abundant transcripts regardless of carbon or nitrogen source. Metabolome analysis yielded 341 metabolites across 16 samples (quadruplicate of each carbon-nitrogen combination; **Table S2**). There were no significant differences in transcript detection among treatments, but significantly fewer metabolites were detected in the methanol-ammonium cultures (**Fig. S1**). Both supervised and unsupervised clustering approaches confirmed that the treatments exhibited distinct transcriptome and metabolome profiles (**Fig. 2, Fig. S2**).

**Fig. 1:**
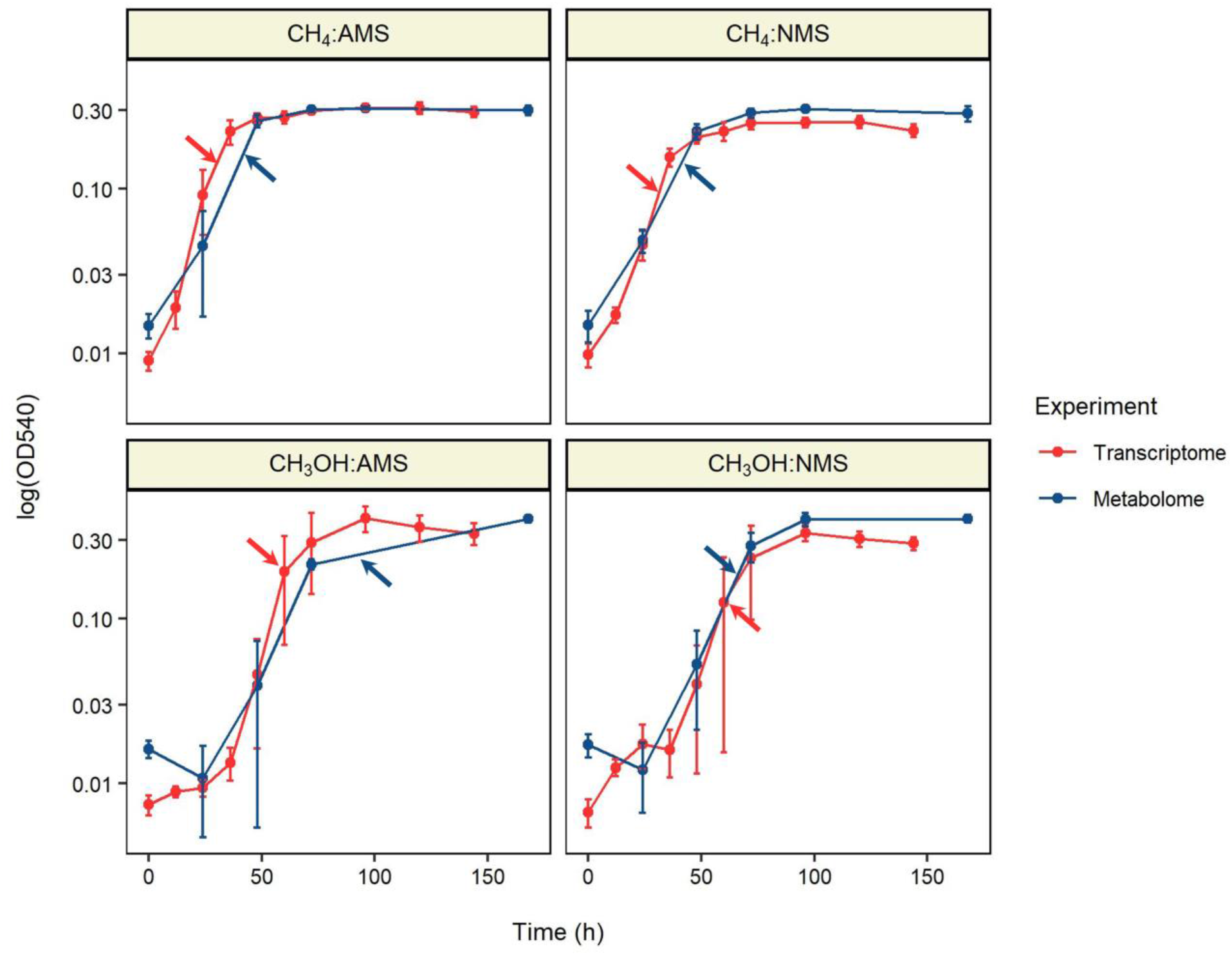
Growth curves for *Methylomicrobium album* BG8 under different media conditions. Growth was measured as the optical density (OD) at 540 nm of cultures of *M. album* BG8. Cultures were provided with either methane or methanol as the carbon source and either ammonium (AMS) or nitrate (NMS) as the nitrogen source. Arrows indicate when cultures were harvested for transcriptomic (red) or metabolomic (blue) analysis.

**Fig. 2:**
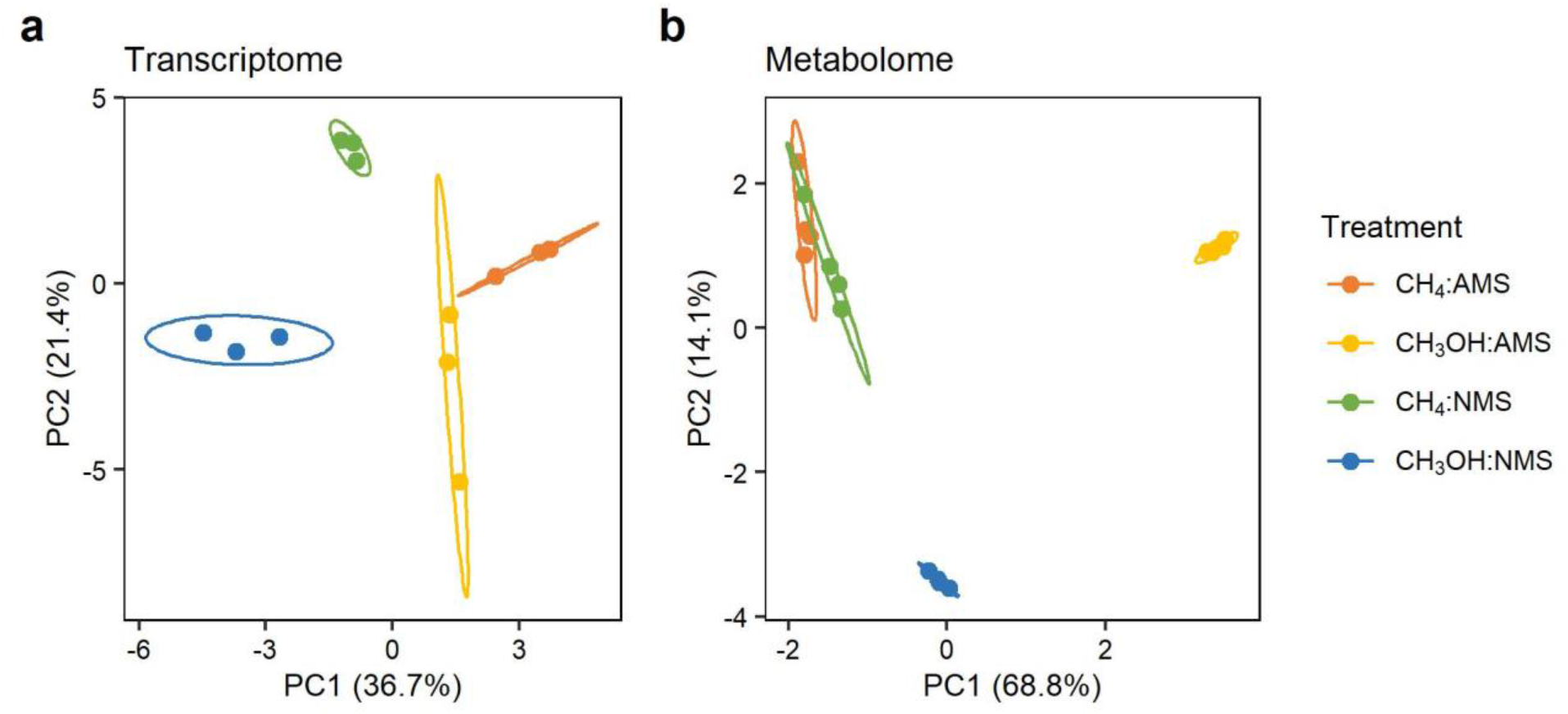
Principal components analysis. Principal components analysis was performed on natural log-transformed transcript RPKM values (**a**) and on natural log-transformed median-scaled metabolite abundances (**b**). Ellipses indicate 95% confidence intervals.

### Carbon source effects when using nitrate

To determine how *M. album* BG8 maintains equivalent growth in methane and methanol, we first compared transcriptomic and metabolomic responses between methane and methanol when cultures were grown using nitrate. In total, 257 genes were significantly differentially expressed between treatments (**Fig. 3a**), and the number of up- and down-regulated genes was equally distributed between methane (n=128) and methanol (n=129). Notably, over 50% of the genes upregulated in methanol had no functional annotation (**Table S1**). Hypergeometric test-based over-representation analysis showed that methanol samples were enriched only for “translation” (J) (**Fig. 3b; Table S3**), with almost all ribosomal proteins showing moderate to significant upregulation in methanol. The stress response sigma factor *rpoE*, carbon storage regulator *csrA*, and other oxidative stress response genes were also upregulated in methanol (**Table S1**). Cells grown in methane were enriched for “cell motility” (N), “inorganic ion transport” (P), and “secondary metabolite biosynthesis” (Q) (**Fig. 3b; Table S3**); representative upregulated genes in these categories included flagellar proteins, metal transporters, and non-ribosomal peptide synthases, respectively (**Table S1**). Genes involved in oxidative phosphorylation, including cytochrome *c* oxidase and NADH ubiquinone oxidoreductase, were also consistently more abundant in methane, though not all these differences were significant (**Table S1**).

**Fig. 3:**
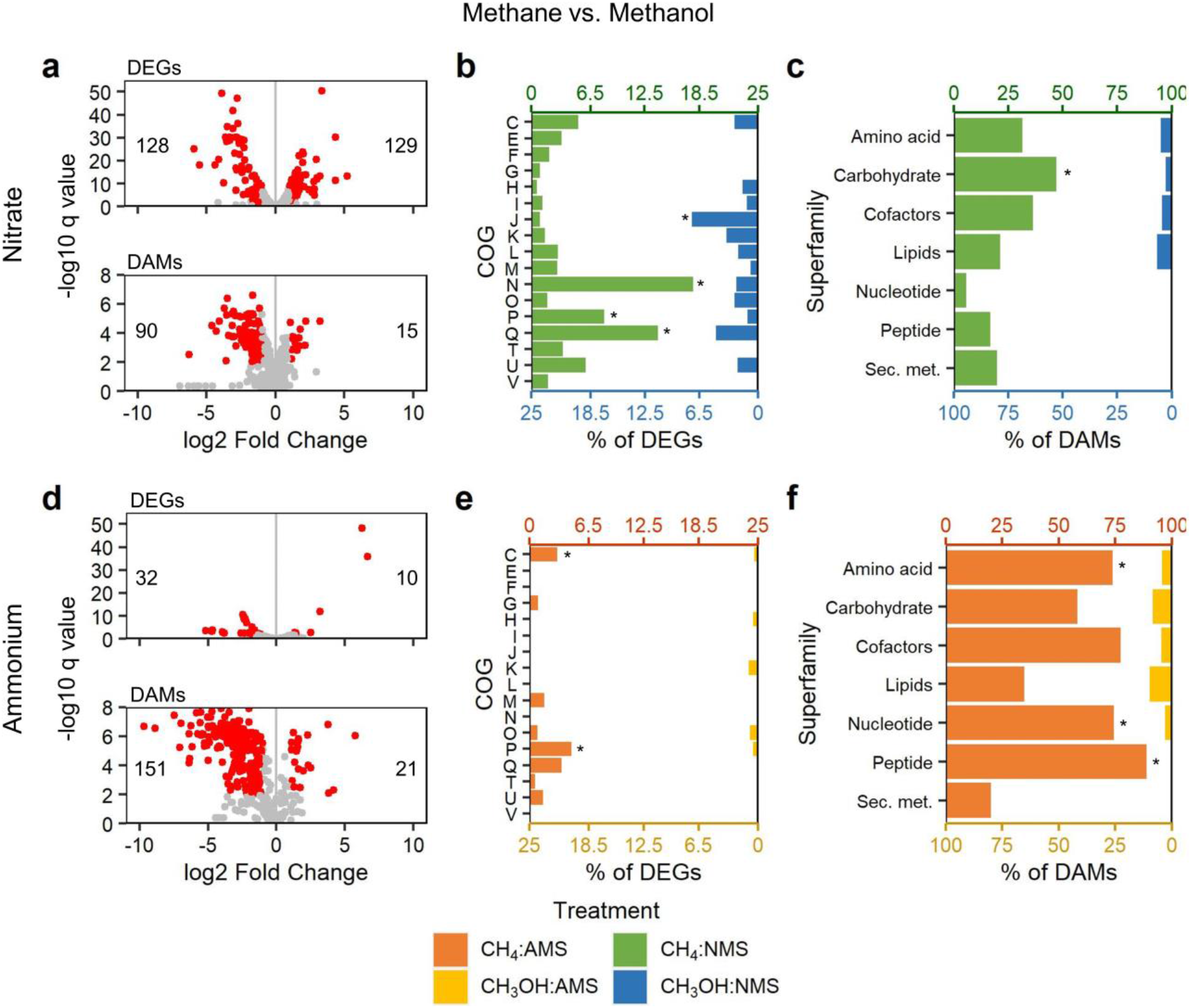
Transcriptomic and metabolomic responses between methane and methanol. Transcriptomic and metabolomic responses to methanol are shown with both nitrate (top panels) and ammonium (bottom panels) as a nitrogen source. (**a, d**) Volcano plots for differentially expressed genes (DEGs) and differentially abundant metabolites (DAMs) between methane and methanol, using methane as a reference. Numbers indicate the number of DEGs or DAMs that were upregulated (on right) and downregulated (on left) when cultures were grown on methanol rather than methane. (**b, e**) DEGs were categorized based on the Clusters of Orthologous Groups (COG) database. For each COG, the number of DEGs is expressed as a percentage of the total number of detected genes in that COG. Significantly over-represented COGs (p < 0.05) are indicated with asterisks. (**c, f**) DAMs were similarly categorized according to their superfamily classification and expressed as a percentage of the total number of detected metabolites in that superfamily. Significant over-representation (p < 0.05) is indicated with asterisks.

Metabolite production was highly favored in methane cultures versus methanol: of the 105 DAMs distinguishing methane-nitrate and methanol-nitrate cultures, 90 DAMs were more abundant in methane (**Fig. 3a**). These metabolites were distributed relatively equally among amino acids, carbohydrates, and cofactors, though only carbohydrates were significantly over-represented (**Fig. 3c; Table S4**). No metabolite superfamilies or subfamilies were significantly over-represented in methanol cultures (**Fig. 3c; Tables S4-5**); the few metabolites that were significantly more abundant in methanol included select fatty acids, phospholipids, lyso-phospholipids, aromatic amino acids, and branched chain amino acids, as well as histidine and histidinol (**Table S2**). Several additional phospholipids were more abundant in methanol cultures, but the fold-changes between treatments did not meet our significance criterion (**Table S2**).

The DEGs and DAMs between methane-nitrate and methanol-nitrate cultures were used for KEGG pathway enrichment analysis to test for differences in general metabolic activity. Significantly affected pathways among the DEGs included flagellar assembly, which was more abundant in methane, and ribosomal biosynthesis, which was more abundant in methanol (**Table S5**). Among the DAMs, significantly over-represented KEGG pathways included select amino acid biosynthesis pathways, which were largely implicated by the increased abundances of glutamate, glutamine, pyruvate, and TCA cycle intermediates in methane cultures (**Table S5**). However, many of the DAMs, including phospholipids and branched chain and aromatic amino acids, were not assigned to any KEGG pathways, limiting the utility of this test.

We manually inspected other pathways and noted that 6-phosphogluconate dehydrogenase and glucose-6-phosphate 1-dehydrogenase, two key enzymes shared by the ED and PP pathways, were among the most highly upregulated DEGs in methanol cultures (**Fig. 4; Table S1**), whereas several of the enzymes and metabolites involved in the EMP glycolytic pathway and the TCA cycle were upregulated in methane cultures (**Fig. 4; Table S1**). Genes encoding glutathione-dependent formaldehyde detoxification to CO2 via S-hydroxymethylglutathione, S-formylglutathione, and formate were also upregulated in methanol cultures (**Fig. 4; Table S1**), whereas all detected gamma-glutamyl amino acids, products of glutathione degradation, were either slightly or significantly more abundant in methane cultures (**Table S2**). **Figure 4** provides an integrated visual representation of the transcriptomic and metabolomic differences between methane-nitrate and methanol-nitrate cultures in the context of their biological pathways and functions.

**Fig. 4:**
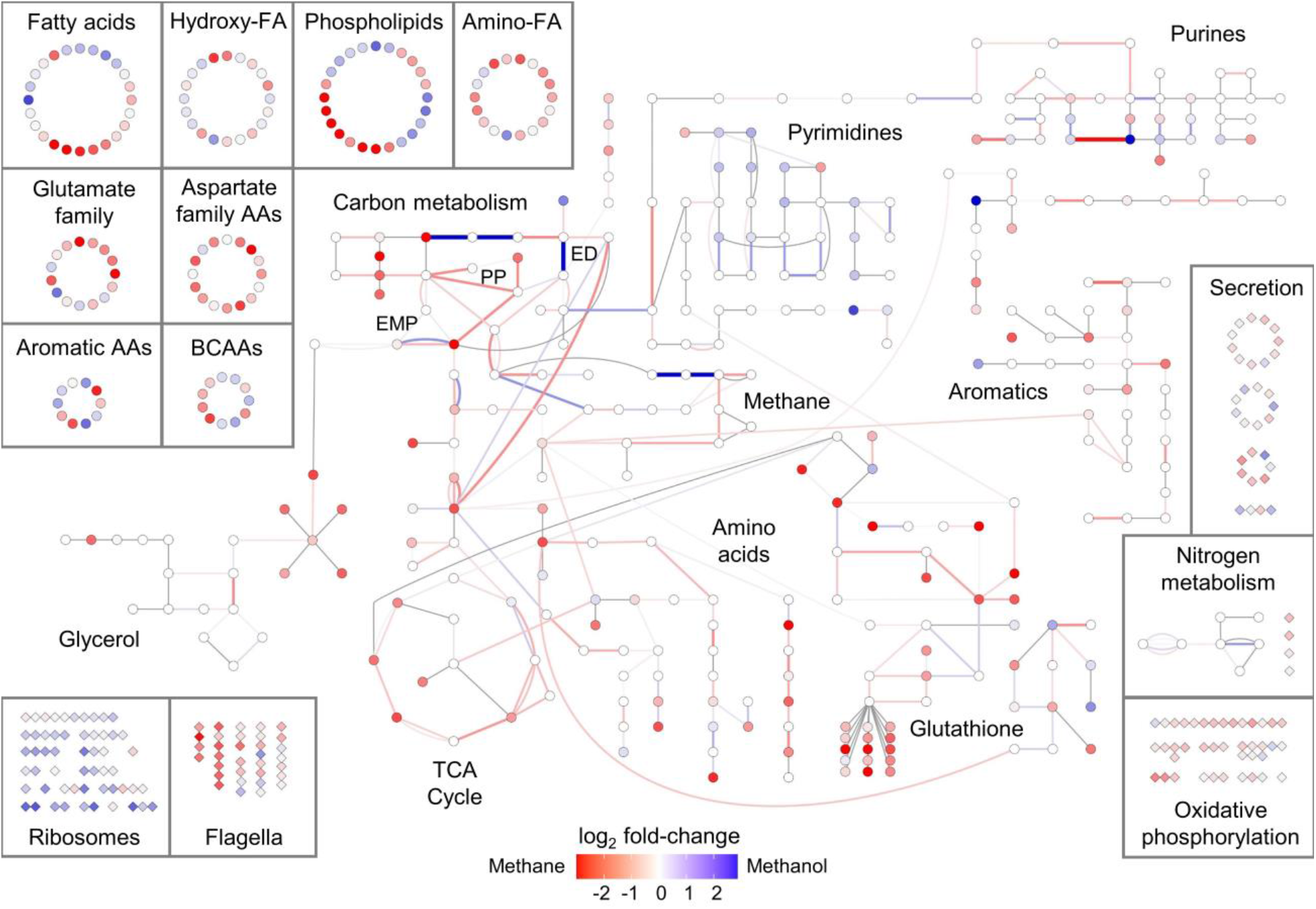
Pathway-level responses to methane and methanol. Central metabolic pathways are presented in the center of the figure. Additional metabolites not involved in central metabolic pathways are shown in the top left, and additional transcripts are shown in the bottom left and right. Transcripts (connections) and metabolites (nodes) are colored based on their log2-transformed fold change between methane and methanol for cultures grown in nitrate; red and blue indicate upregulation in methane or methanol, respectively. See **Fig. S3** for the same figure reproduced for cultures grown in ammonium. Abbreviations: FA, fatty acids; AA, amino acids; BCAA, branched chain amino acids; ED, Entner-Doudoroff; EMP, Embden-Meyerhof-Parnas; PP, pentose phosphate; TCA, tricarboxylic acid.

### Carbon source effects when using ammonium

We then tested whether the observed transcriptomic and metabolomic responses to growth on methane versus methanol were consistent when ammonium was substituted for nitrate as the nitrogen source. The broad trends in the transcriptome were largely the same, with methanol-grown cultures showing upregulation of all ribosomal proteins and methane-grown cultures showing upregulation of flagellar proteins, oxidative phosphorylation enzymes, metal transporters, and, to a lesser extent, stress response genes (**Table S1**). However, the effect size of many of these differences between treatments was smaller in ammonium: only 42 genes were identified as DEGs based on our criteria for significance, six-fold fewer than in nitrate (**Fig. 3d**). The only exception was genes for oxidative phosphorylation, which were upregulated in methane and had a higher fold-change in ammonium, resulting in significant over-representation at both the COG (**Fig. 3e; Table S3**) and KEGG (**Table S5**) levels.

In contrast, the effect size of metabolomic differences was two-fold larger in ammonium than nitrate (**Fig. 3d, f**), even when excluding metabolites that were not detected in methanol-ammonium cultures. The broad trends remained similar, however, with the 21 methanol-enriched DAMs encompassing select fatty acids, phospholipids, branched chain amino acids, and histidinol. One notable difference between carbon source effects in ammonium and nitrate was that seduheptulose-7-phosphate, a key PP pathway intermediate, was significantly more abundant in methanol-ammonium relative to methane-ammonium cultures (**Table S2**), though methane cultures still preferred the EMP pathway whereas methanol cultures favored the ED and PP pathways. These changes are visually represented in **Fig. S3**.

### Nitrogen source effects

Lastly, we evaluated how *M. album* BG8 responds to nitrate and ammonium, and whether these results were consistent in both methane and methanol. In methane cultures, there were 106 DEGs and 25 DAMs between ammonium and nitrate treatments (**Fig. 5a**). Nitrate-grown cultures were significantly enriched for genes associated with “inorganic ion transport and metabolism” (COG P) (**Fig. 5b**) and the KEGG pathway “nitrogen metabolism” (**Table S5**). These genes primarily included ABC transporters involved in acquiring extracellular nitrate as well as the nitrate and nitrite reductase genes responsible for converting nitrate into bioavailable ammonium (**Table S1**). Glutamine synthetase, glutamine, select sphingolipids, and genes for iron transport were also upregulated in nitrate cultures (**Table S2**). Ammonium-grown cultures, in contrast, were enriched for “energy production and conversion” (COG C) and the KEGG pathway “oxidative phosphorylation” (**Fig. 5e, Table S5**), and generally exhibited higher levels of transcripts and metabolites associated with basic metabolism, including the TCA cycle and nucleotide biosynthesis (**Fig. 6**). Transcripts for hydroxylamine dehydrogenase, *haoAB*, and several efflux pumps were also upregulated in ammonium cultures (**Fig. 6; Fig. S4; Table S1**).

**Fig. 5:**
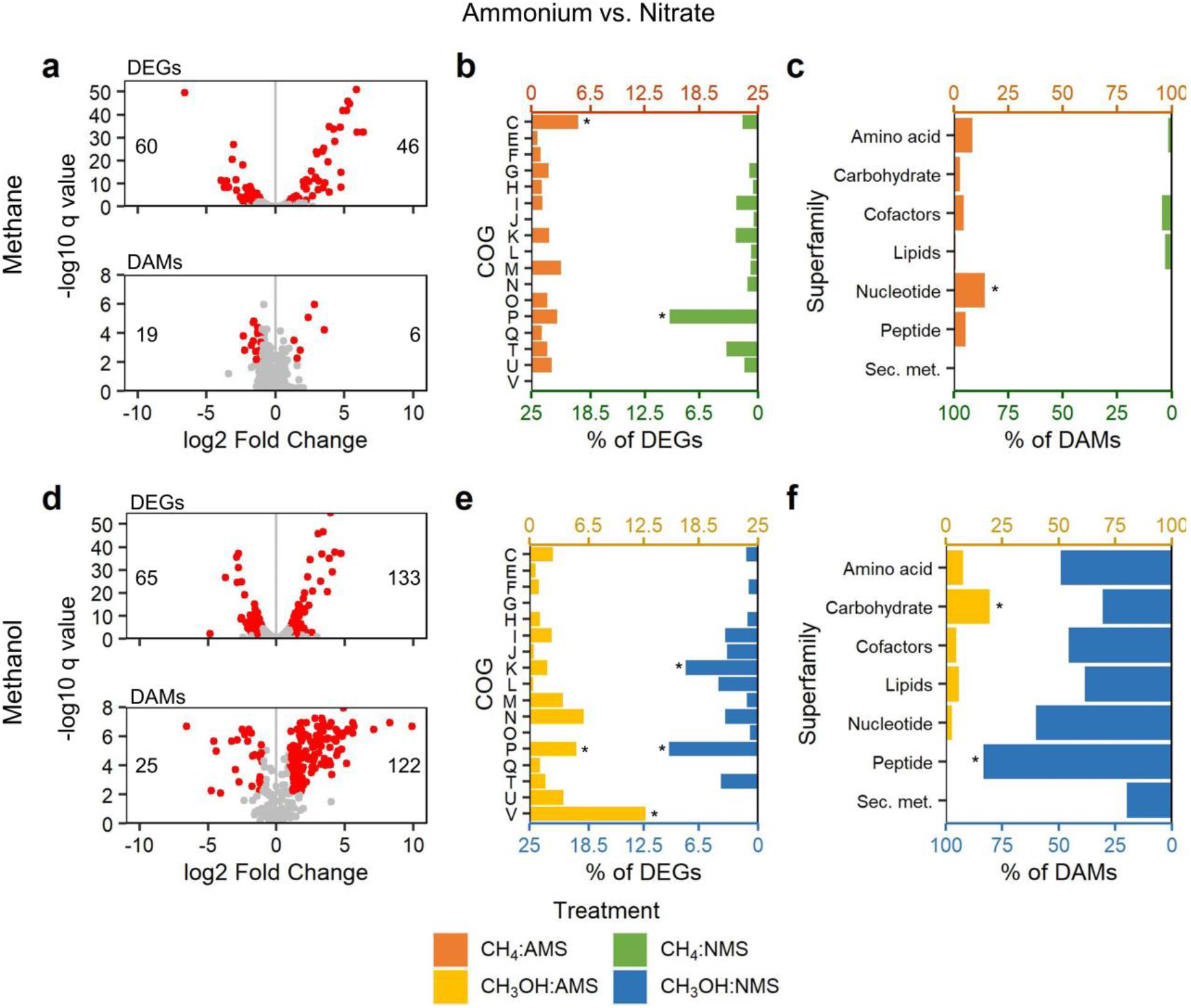
Transcriptomic and metabolomic responses between ammonium and nitrate. Transcriptomic and metabolomic responses to nitrogen source are shown with both methane (top panels) and methanol (bottom panels) as a carbon source. Refer to Figure 3 for a complete description of the panels. Fold changes are expressed relative to nitrate, so that numbers in (**a**) and (**d**) indicate the number of DEGs or DAMs that were upregulated (on right) and downregulated (on left) when cultures were grown on ammonium rather than nitrate.

**Fig. 6:**
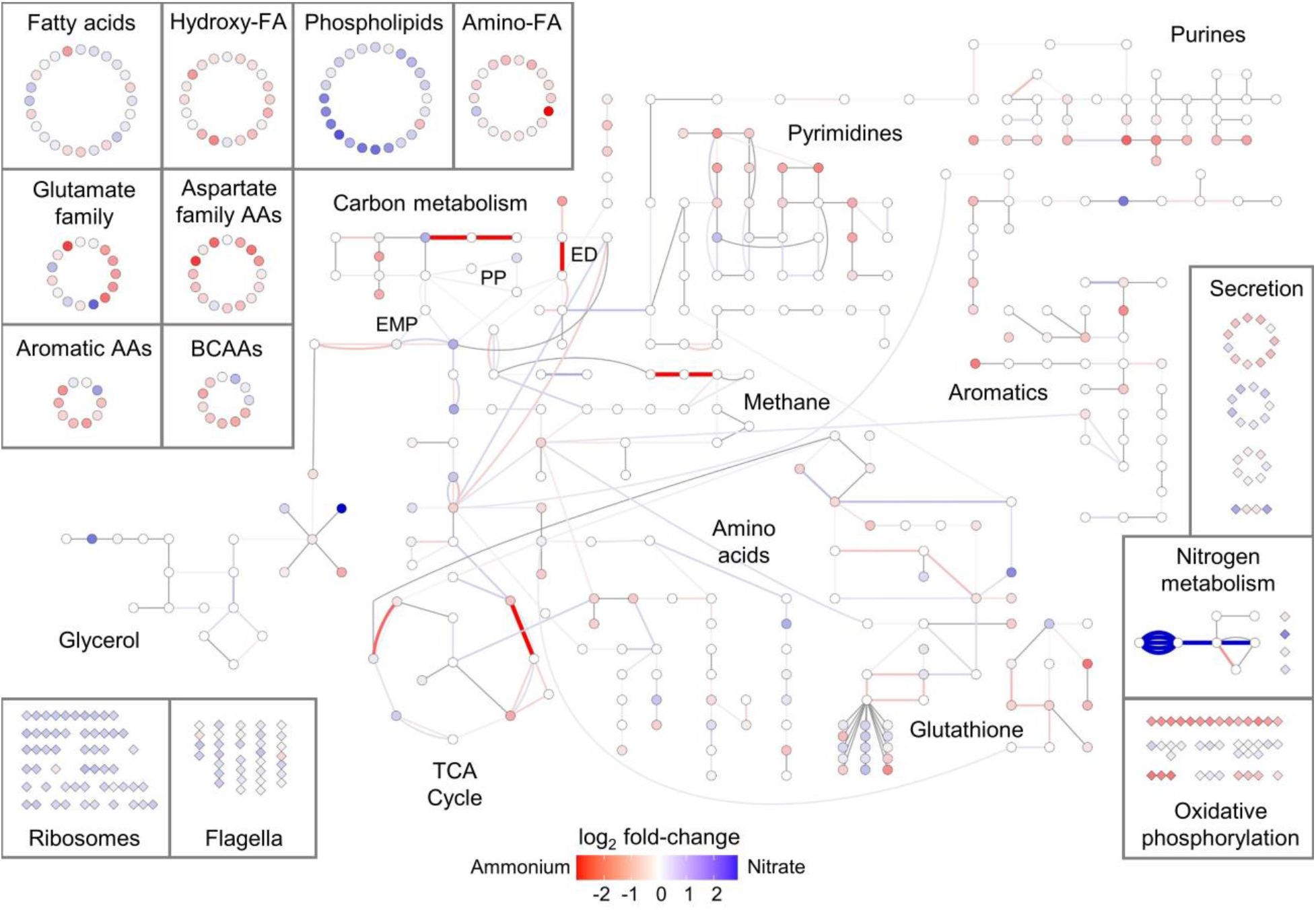
Pathway-level responses to ammonium and nitrate. Central metabolic pathways are presented in the center of the figure. Additional metabolites not involved in central metabolic pathways are shown in the top left, and additional transcripts are shown in the bottom left and right. Transcripts (connections) and metabolites (nodes) are colored based on their log2-transformed fold change between ammonium and nitrate for cultures grown in methane; red and blue indicate upregulation in ammonium or nitrate, respectively. See **Fig. S4** for the same figure reproduced for cultures grown in methanol. Abbreviations: FA, fatty acids; AA, amino acids; BCAA, branched chain amino acids; ED, Entner-Doudoroff; EMP, Embden-Meyerhof-Parnas; PP, pentose phosphate; TCA, tricarboxylic acid.

The nitrogen source response was stronger in methanol cultures, with two-fold more DEGs (n=198) and six-fold more DAMs (n=147) than in methane cultures (**Fig. 5**). Nitrate assimilation and conversion genes were significantly upregulated in nitrate cultures, whereas *haoAB* genes were significantly upregulated in ammonium cultures (**Table S1; Fig. S4**), as before, but many other trends became less clear for cells grown on methanol, even when considering only the metabolites that were detected in both treatments. General transcriptional regulators (COG K) were significantly over-represented in nitrate cultures, as were dipeptides, whereas drug efflux pumps (COGs P and V) were significantly over-represented in ammonium cultures (**Fig. 5e, f**). Basic metabolic activity appeared to be higher in nitrate rather than ammonium, with most nucleotides and amino acids accumulating at higher abundances in nitrate-grown cultures (**Fig. S4**). Central carbon pathways were the only exception to this trend, with ammonium-grown cultures showing general upregulation of the genes and metabolites involved in central carbon metabolism.

## DISCUSSION

An understanding of the growth and physiology of specific methanotroph strains in response to carbon-nitrogen source is important for formulating optimal growth media and for scaling up their growth in an industrial context. Using a global transcriptomic and metabolomic approach, we evaluated how *M. album* BG8 maintains equivalent growth rates when growing on either methane or methanol, and whether its responses to either ammonium or nitrate are consistent between the two carbon sources. We found evidence that, with nitrate as the nitrogen source, *M. album* BG8 maintains growth in methanol by increasing flux through the ED and PP pathways (6), synthesizing additional ribosomes, restructuring its phospholipid membrane, and upregulating other stress response signatures while downregulating almost all other metabolic activity. These responses were consistent when cultures were provided with ammonium, which has not been tested in other studies of methanotroph metabolism growing on methanol (5, 6, 12). In addition, we found that the differences between ammonium- and nitrate-grown cultures were unexpectedly magnified in methanol, though the affected pathways and metabolisms were largely the same. Several of the physiological responses we observed could only be detected using the global multi-omics approach used here and have not been described in previous studies of central metabolic pathways in other gammaproteobacterial methanotrophs (5, 6, 12, 15).

We identified five major differences distinguishing growth on methanol from growth on methane, most of which presumably stem from the effects of methanol on redox chemistry and the general cellular stress of growing on an organic solvent. First, methanol-grown *M. album* BG8 appeared to prefer the ED and PP pathways for sugar catabolism. In *M. buryatense* 5GB1, the ED pathway is essential for cellular growth, though the reasons for its importance remain unclear (18). Increased carbon flux through the ED pathway for *M. buryatense* 5GB1 growing on methanol has been attributed to the lower rate of ATP production from this pathway, as the ATP production power in methanol can instead be provided by MDH-derived electrons that are not needed for methane oxidation (6). Our observation that the NADH oxidoreductases are downregulated in methanol cultures provides further support for this hypothesis; in methanol, MDH-derived electrons can be directly transferred to cytochromes to fuel the electron transport chain (25), reducing the need for the electron-harvesting NADH oxidoreductase complex to power ATP production. There is less evidence for the importance of the PP pathway in methanotrophs, though increased flux through the PP pathway has been shown to satisfy the higher demand for NADPH production in response to oxidative stress in *M. alcaliphilum* 20Z (26). Other gammaproteobacterial methanotrophs do not show the same preference for the ED or PP pathways when growing on methanol (5, 17), suggesting that some methanol adaptations may be strain-specific, but further investigation of sugar catabolism pathway flux using more targeted analysis will be needed to confirm carbon pathway preferences in *M. album* BG8.

Secondly, we found that growth on methanol induced dramatic and consistent upregulation of translational machinery, including all ribosomal proteins. Bacterial growth is controlled by the rate of protein synthesis, which is in turn dictated by the number of actively translating ribosomes and the translational elongation rate (27). Previous studies have shown that *E. coli* responds to stressors that lower the translation elongation rate, such as reactive oxygen species, by increasing ribosome production, while the opposite is true for stressors that reduce ribosome production (27). Because ribosomes are rate-limited (28), the production of additional ribosomes by methanol-grown *M. album* BG8 may therefore be a strategy for maintaining the same growth rate while growing on a toxic substrate. We hypothesize that the longer lag phase described for methanol-grown methanotrophs, which was also observed for *M. album* BG8, may be caused by this requirement for additional ribosomes to be produced when growing on methanol, as cultures cannot begin logarithmic growth until they have produced a sufficient number of ribosomes (28–30).

Thirdly, we observed significant alterations in fatty acid and phospholipid production that may reflect membrane adaptation to growth on methanol, an organic solvent. Organic solvents are generally toxic to bacteria because they compromise cell membrane integrity (31); however, several species of Gram-negative bacteria are able to thrive in the presence of solvents by altering cell membrane composition (32), upregulating solvent efflux pumps in the outer membrane (33), and increasing the rate of solvent biotransformation (34). We did not find evidence for the latter two mechanisms acting in *M. album* BG8 growing on methanol, but the significant differences in fatty acid and phospholipid abundances that we observed likely indicate some degree of membrane remodeling. Although we expected unsaturated fatty acids would be more abundant in methanol cultures, reflecting the increased membrane fluidity required to uptake methanol from the culture media (35), there were no clear trends in the abundances of saturated or unsaturated lipids. We hypothesize that this is driven by the need to balance methanol assimilation with the need to ensure membrane integrity when exposed to a solvent, though further work will be needed to test this hypothesis. Growth on methanol has also been shown to decrease fatty acid methyl ester abundance in *M. album* BG8 due to the reduced need to synthesize intracytoplasmic membranes for methane oxidation (13). The metabolomics approach used in this study did not allow us to quantify lipid abundances, but our results nonetheless provide evidence for alterations in lipid composition as a mechanism for methanol tolerance.

Fourthly, we observed other general responses that indicated the presence of methanol led to a more stressful environment for the cells. Methanol-grown cultures had two-fold higher abundances of the envelope stress response sigma factor *rpoE* and the carbon storage regulator *csrA*, which are required for maintaining periplasmic and outer membrane integrity (36) and adjusting central carbon metabolism to stress conditions (37), respectively. We also observed upregulation of several oxidative stress responses described for *Methylomonas* sp. DH-1 grown on methanol (17), including hopanoid biosynthesis, peroxiredoxin, superoxide dismutase, and thioredoxin, though we did not observe the corresponding changes in the *oxyR* or *soxS* regulons. Select branched-chain and aromatic amino acids, which are associated with adaptation to nutrient limitation (4, 38), were also upregulated in methanol cultures. Aside from these few exceptions, most central metabolic pathways were downregulated in the presence of methanol: methanol-grown *M. album* BG8 accumulated fewer carbohydrates, TCA cycle intermediates, and nucleotides, and downregulated several of the genes involved in those pathways. These results align with the well-established observation that stress reduces global metabolic activity (39), as energy stores are overwhelmingly directed towards growth, leaving little room for the production of other metabolites or energy-intensive cellular accessories such as flagella (40, 41).

Finally, *M. album* BG8 cultures produced significantly fewer gamma-glutamyl amino acids, 5-oxoproline, and glutamate when grown on methanol, indicating the upregulation of GSH-dependent formaldehyde detoxification in methanol cultures and the upregulation of glutathione (GSH) degradation in methane cultures. One of the three formaldehyde detoxification strategies in bacteria is GSH-dependent (42). During this response, GSH is condensed with formaldehyde either spontaneously or via formaldehyde-activated enzyme to yield S-hydroxymethylglutathione (HMGS), which is subsequently converted to formate through a multi-step reaction that releases GSH. The regenerated GSH can then be used to detoxify another formaldehyde molecule (42). We noted that several of the genes involved in these reactions were also upregulated in methanol cultures. However, if the cell does not experience formaldehyde toxicity, unused GSH can instead be degraded; during this process, gamma-glutamyl transferase breaks down GSH and transfers the gamma-glutamyl group to an amino acid. In later steps, these gamma-glutamyl acids can be converted to 5-oxoproline, which is used to produce glutamic acid (43). The methanol-induced upregulation of genes involved in GSH-dependent formaldehyde detoxification, and the lower abundance of the byproducts of GSH degradation, collectively suggest that *M. album* BG8 cultures grown in methanol upregulate the GSH-dependent formaldehyde detoxification system to overcome formaldehyde toxicity. Although we lack clear evidence for this, it is possible that *M. album* BG8 also uses a partial serine cycle to fuel tetrahydrofolate-based formaldehyde detoxification (3, 42), which would further enable growth on methanol.

These five adaptations to methanol (altered carbon flux, ribosomal biogenesis, phospholipid replacement, stress responses, and formaldehyde detoxification) were largely conserved when cultures were grown in ammonium: methanol-grown cultures again exhibited significant upregulation of translation machinery, downregulation of oxidative phosphorylation enzymes, preference for the ED and PP pathways, and decreased general metabolic activity. However, there was no significant change in the expression of *rpoE*, superoxide dismutase, or thioredoxin, though other stress response genes remained upregulated in methanol cultures. The accumulation of sedoheptulose-7-phosphate suggested that the PP pathway was slightly preferred over the ED pathway in methanol-ammonium cultures, but more detailed carbon flux analysis would be needed to confirm this hypothesis. Trends in metabolite production when growing on methanol were less clear when ammonium was the nitrogen source, partially due to the low rate of metabolite detection in methanol-ammonium cultures, but we noted that several phospholipids were still significantly differentially abundant between methane and methanol, again suggesting some degree of membrane remodeling. Methanol-ammonium cultures were harvested slightly later than the other treatments (see **Fig. 1**), which may explain some of this variation in metabolite detection.

We additionally examined whether growth on methanol alters the response to nitrogen source, as competitive inhibition of MMO by ammonium would presumably not play a role when the bacteria are not actively oxidizing methane. Although the magnitude of the nitrogen source response was notably larger in methanol cultures, overall trends were largely consistent with previous studies that indicate a hydroxylamine detoxification response in ammonium (20, 22). In both methane and methanol cultures, growth with nitrate expectedly led to the consistent upregulation of all nitrate assimilation and ammonification genes, which would be required to convert nitrogen into bioavailable ammonium. In contrast, the hydroxylamine dehydrogenase genes *haoAB* were upregulated in ammonium cultures, which would allow *M. album* BG8 to rapidly metabolize the hydroxylamine produced by ammonia oxidation by MMO (22). This ammonia-induced upregulation of *haoAB* was preserved even for growth on methanol. Nitrate also led to increased production of lipids, ribosomes, and flagella, whereas ammonium led to the upregulation of oxidative phosphorylation, central carbon pathways, and the TCA cycle. This focus on central metabolic activities for ammonium may explain why ammonium often enriches for gammaproteobacterial methanotrophs (44, 45).

As shown in this study, a global transcriptomic and metabolomic approach provides information on physiological adaptations to nutrient sources that cannot be detected using more focused approaches like ^13^C tracer analysis, which is better for quantifying metabolic flux through specific pathways. It has also been argued that methanotrophs exhibit little in the form of transcriptomic responses and that the proteome or metabolome are therefore a better indicator of substrate-based metabolic responses (6); however, several foci of transcriptional regulation in this study, including ribosome biogenesis, flagellar synthesis, and oxidative phosphorylation enzymes, do not produce detectable metabolites and would not have been identified in a metabolomic study alone. Our results are admittedly limited by the low detection of metabolites in methanol-ammonium cultures and the inability to quantify metabolite abundances in a global metabolomics analysis. Even so, our results implicate several nutrient-based response strategies that have not previously been described for methanotrophs growing with different carbon-nitrogen combinations and provide significant insight and valuable directions for future research on methanotroph physiology and the biotechnological optimization of these species.

## MATERIALS AND METHODS

### Culture growth

We used a two-by-two factorial design, with four different growth conditions derived from the combinations of two carbon sources (methane or methanol) and two nitrogen sources (ammonium or nitrate). Separate cultures of *M. album* BG8 were grown for transcriptomic and metabolomic analyses, and cultures were passaged at least once under experimental conditions before being grown for each experiment. All cultures were grown in 250-ml Wheaton media bottles filled with 100 ml of medium and closed with butyl-rubber septa caps. Cultures were grown either in ammonium mineral salts (AMS) or nitrate mineral salts (NMS) media (46) buffered to pH 6.8 using 1.5 ml phosphate buffer.

For cultures grown with methane, 50 ml of gas headspace was removed from the culture bottle and 2.5 mmol of methane was injected through a 0.22 μm filter-fitted syringe. For cultures grown in methanol, pure HPLC-grade methanol was added to the culture media to a final concentration of 2.5 mmol for the transcriptome cultures and 1 mmol for the metabolome cultures. *M. album* BG8 exhibits similar growth rates at both 1 mmol and 2.5 mmol methanol ((3), see **Fig. 1**), so we expected comparable results between the transcriptome and metabolome experiments. Cultures designated for transcriptome analysis were grown in triplicate and cultures for metabolome analysis were grown in quadruplicate. All cultures were incubated at 30°C with shaking at 150 rpm.

Culture growth was monitored by measuring the optical density of the medium at 540 nm using a 48-well microplate reader (Multiskan Spectrum, Thermo Scientific). Culture purity was assessed via phase-contrast microscopy and plating on TSA/nutrient agar, where colony growth would indicate contamination. Cultures were harvested during logarithmic growth after at least three doublings had occurred (**Fig. 1**), with metabolome cultures harvested slightly later than transcriptome cultures. The delayed metabolite sampling allowed for the effects of transcriptional regulation to be observed at the metabolomic level while still ensuring that all samples were collected during logarithmic growth.

### Transcriptome analysis

Total RNA was extracted from harvested cultures using the Masterpure™ Complete DNA and RNA Kit (Lucigen Corporation, Middleton, Wisconsin). RNA was subsequently purified using the Zymo RNA Clean and Concentrator™ Kit (Zymo Research, Irvine, CA). We assessed the quality and purity of extracted RNA samples using NanoDrop spectrophotometry and an Agilent 2100 Bioanalyzer. High-quality RNA was sent to the *Centre d’Expertise et de Services* at Génome Québec (Montreal, Québec, Canada) for library preparation and sequencing. Ribosomal RNA was depleted from 250 ng of total RNA using the QIAseq FastSelect Kit (Qiagen). RNA was then reverse transcribed using the NEBNext RNA First Strand Synthesis and NEBNext Ultra Directional RNA Second Strand Synthesis Modules (New England BioLabs). The remaining steps of library preparation were done using the NEBNExt Ultra II DNA Library Prep Kit for Illumina (New England BioLabs). Sequencing was performed on an Illumina HiSeq 4000 with 100 bp paired-end reads.

Raw paired-end RNASeq reads were quality-filtered using the default filtering parameters in Trimmomatic 0.39 (47). High-quality reads were mapped to the published *M. album* BG8 genome (48) using BowTie2 2.4.1 (49), and the number of reads mapped to each gene was calculated using HTSeq 0.11.1 (50). To facilitate downstream analyses of functional differences among treatments, we mapped each transcript to functional identifiers in the Clusters of Orthologous Groups (COG) and Kyoto Encyclopedia of Genes and Genomes (KEGG) databases using eggNOG (51) and blastKOALA (52), respectively. For downstream analyses requiring normalized count data, we converted raw HTSeq counts to transcripts per million (TPM).

### Metabolome analysis

Frozen cell pellets were processed by Metabolon Inc. (Durham, NC, USA) for global metabolomics analysis using the Metabolon HD4 platform. In brief, metabolites were extracted using methanol with vigorous shaking and then recovered by centrifugation. Methanol was removed using a TurboVap^®^ (Zymark). The resulting extracts were stored under nitrogen overnight before being analyzed using four independent procedures: 1) reverse-phase ultra-high performance liquid chromatography-tandem mass spectroscopy (RP/UPLC-MS/MS) with positive ion mode electrospray ionization (ESI), chromatographically optimized for hydrophilic compounds; 2) the same procedure optimized for hydrophilic compounds; 3) RP/UPLC-MS/MS with negative ion mode ESI; and 4) hydrophilic interaction chromatography (HILIC)/UPLC-MS/MS with negative ion mode ESI. All analysis procedures were performed alongside multiple quality control samples as outlined in Metabolon’s standard protocol (www.metabolon.com).

Metabolites were identified by comparing the retention time, mass-to-charge ratio, and chromatographic data for each compound to a database of known standards. This process additionally classified metabolites into seven main superfamilies based on their molecular structure (amino acids, carbohydrates, cofactors and electron carriers, lipids, nucleotides, peptides, and secondary metabolites). Metabolite abundances were quantified as area-under-the-curve detector ion counts. If a metabolite was detected in some samples, but not all, missing values were assumed to be lower than the detection threshold for the analysis platform and were therefore imputed as one-half the lowest detected abundance of that metabolite. For downstream analyses requiring normalized data, we divided raw area-under-the-curve values by the median value for each metabolite prior to minimum value imputation.

### Statistical analysis

All downstream analyses were performed in R 3.6.2 (53). For both transcriptome and metabolome data, we first tested for broad differences among treatments by performing principal components analysis (PCA) and sparse partial least squares discriminant analysis (sPLS-DA). These analyses were performed on natural log-transformed transcripts per million (TPM) values and natural log-transformed median-scaled metabolite abundances using the ‘rda’ function in the R package *vegan* (54) and the ‘splsda’ function in the package *mixOmics* (55). We used an ANOVA with Tukey’s *post hoc* test to evaluate if there were differences in transcript or metabolite detection rates among treatments.

We then evaluated the effects of carbon and nitrogen sources on metabolic activity in *M. album* BG8 by testing for differentially expressed genes (DEGs) and differentially abundant metabolites (DAMs). DEGs and DAMs were identified for each of the four pairwise comparisons in our two-by-two design. All differential abundance analyses were performed on raw counts or abundances using *edgeR* (56) for transcriptome data and the Student’s t-test for metabolome data. P-values were adjusted using the Benjamini-Hocherg (FDR) correction, and transcripts or metabolites with a log_2_-transformed fold change > |1| and FDR < 0.01 were considered significant. Due to the significantly lower metabolite detection rate in methanol-ammonium (see Results and **Fig. S1**), we chose to remove undetected metabolites from all pairwise DAM comparisons that included this treatment, which resolved the large number of interaction effects originally observed in the metabolome data. We additionally confirmed the differential abundance results for this treatment by comparing to results obtained using a separate procedure in which within-sample metabolite abundances were normalized prior to differential abundance testing (see **Supplementary Methods**).

To determine whether any physiological functions or metabolic pathways were preferentially enriched in the DEGs and DAMs associated with each treatment comparison, we categorized DEGs and DAMs based on COGs and metabolite superfamilies, respectively. We tested for significant over-representation of COGs and of metabolite superfamilies and subfamilies using Fisher’s exact test, with a significance threshold of FDR-adjusted p < 0.05. Over-representation analyses were repeated at the pathway level after assigning transcripts and metabolites to their corresponding pathways in the KEGG pathway database. We used the package *clusterProfiler* (57) to test for pathway over-representation in the transcriptome data and *MetaboAnalyst* (58) for the metabolome data, with a significance threshold of FDR-adjusted p < 0.05. Transcripts and metabolites that did not have a corresponding KEGG ID were excluded from over-representation analyses.

## Data availability

Raw transcriptome reads have been deposited in the NCBI Short Read Archive (SRA) under the accession number PRJNA698057. Metabolome data has been deposited in the MetaboLights database under the accession number MTBLS2127 (www.ebi.ac.uk/metabolights/MTBLS2127). The R code and workspace required to reproduce all analyses can be found in the GitHub repository https://github.com/sasugden/BG8_multi_omics.

## DECLARATIONS

### Author contributions

Conceptualization, S.S., M.L., D.S. and L.Y.S.; data curation, S.S.; formal analysis, S.S.; funding acquisition, D.S. and L.Y.S.; investigation, S.S. and M.L.; project administration, D.S. and L.Y.S.; resources, D.S. and L.Y.S.; software, S.S.; supervision, L.Y.S.; visualization, S.S.; writing—original draft preparation, S.S.; writing—review and editing, S.S., M.L., D.S., and L.Y.S. All authors have read and agreed to the published version of the manuscript.

## Acknowledgements

The authors would like to thank Phillip Sun, Melissa Harington, and Mariah Hermary for technical assistance with culturing and RNA extraction. Danny Alexander and Metabolon assisted with the analysis and interpretation of the metabolomics data.

